# Elemental stoichiometry and insect chill tolerance: Evolved and plastic changes in organismal Na^+^ and K^+^ content in *Drosophila*

**DOI:** 10.1101/2022.08.18.504319

**Authors:** Sarah C. Chalmer, Seth M. Rudman, Mads K. Andersen, Paul Schmidt, Heath A. MacMillan

## Abstract

Acclimation and evolutionary adaptation can produce phenotypic change that allows organisms to cope with challenges like those associated with climate change. Determining the relative contributions of acclimation and adaptation is of central importance to understanding animal responses to change. Rates of evolution have traditionally been considered slow relative to ecological processes that shape biodiversity. Many organisms nonetheless show patterns of spatial genetic variation suggestive of adaptation and some evidence is emerging that adaptation can act sufficiently fast to allow phenotypic tracking in response to environmental change (‘adaptive tracking’). In *Drosophila*, both plastic and evolved differences in chill tolerance are associated with ionoregulation. Here we combine acclimation, latitudinal field collections, and a replicated field experiment to assess the effects of acclimation and adaptation on chill coma recovery and elemental (Na and K) stoichiometry in both sexes of *Drosophila melanogaster*. Acclimation and spatial adaptation both shape chill coma recovery, with acclimation producing the greatest magnitude response. Leveraging knowledge on the physiological mechanisms that underlie variation in chill tolerance traits, we find that the relationship between K content and chill tolerance differs among flies acclimated vs. adapted to cold. Taken together, these data reinforce the importance of acclimation in responses to abiotic challenges and illustrate that the mechanisms of phenotypic change can differ between acclimation and basal tolerance adaptation.

## Introduction

Understanding the pace, magnitude, and mechanisms of organismal responses to changing climates is a major goal in biology with profound implications for the conservation of biodiversity (Urban 2015). Organismal traits and their underlying physiology can be critical determinants of both the distribution of biodiversity and population growth rates. Traditionally, phenotypic responses to rapidly changing environments have focused on measuring phenotypic plasticity, as these plastic shifts occur within a generation (Pigliucci 2001). Many organisms exhibit profound phenotypic plasticity, including numerous cases of acclimation in response to abiotic change (Nylin and Gotthard 1998; Ghalambor et al. 2007). Similarly, there are many examples of spatial intraspecific genetic variation indicative of adaptation to abiotic factors that vary over space (Hoffmann et al. 2002; Rajpurohit et al. 2018; des Roches et al. 2020). In the past few decades there has been a growing recognition that populations can also exhibit evolutionary phenotypic change in response to environmental shifts over ecological timescales (Reznick et al. 1990; Hendry and Kinnison 1999; Hairston et al. 2005). This includes evidence that selection can be sufficiently strong to drive adaptive phenotypic tracking over contemporary timescales (Barrett et al. 2008; Behrman et al. 2015; Rennison et al. 2019). Yet, the pace of evolutionary adaptation in response to abrupt changes in climate and its importance in driving phenotypic shifts relative to acclimation is still largely unknown.

Documenting adaptation in response to changes in climate requires demonstration that phenotypic shifts are genetically based (often determined using common garden experiments) and that the phenotypic change leads to an increase in fitness (Merilä and Hendry 2014). Links between phenotypic change and fitness change are difficult to empirically measure, but cases of parallel phenotypic evolution across replicate populations or environmental gradients present strong evidence for adaptation (Losos 1992; Schluter and Nagel 1995). Most of what is known about adaptation comes from examining phenotypic differences among divergent populations or species using phylogenetic comparative methods or comparing spatially separated populations (Futuyma and Kirkpatrick 2005; Kellermann et al. 2012). These approaches are powerful and essential when adaptation occurs too slowly to observe directly, but they don’t allow for precise estimates of evolutionary rates that are needed to assess responses to rapidly changing environments (Gingerich 1993). If adaptation occurs rapidly, it is possible to use longitudinal studies and manipulative experiments to directly measure the extent and pace of adaptation in response to environmental change, termed adaptive tracking (Grant and Grant 1995; Botero et al. 2015; Rudman et al. 2022). Combined with assessments of adaptation arising across populations, which results from dozens to thousands of generations of evolutionary divergence, direct observation of adaptive tracking provides the temporal resolution needed to compare the impact of evolution and phenotypic plasticity on rapid trait change.

With climates changing rapidly due to anthropogenic disturbance it is particularly important to understand the phenotypic responses, both plastic and evolved, to physiologically challenging temperatures. In insects, a primary physiological challenge linked to distribution is chill susceptibility. Chill-susceptible insects are those that can withstand cool temperatures above their freezing point for an extended period but accumulate chilling injuries while doing so (Bale 1996; Andersen et al. 2015; Overgaard and MacMillan 2017). A primary mechanism of these injuries is the disruption of organismal ion homeostasis leading to cell death. Exposure to cold temperatures is associated with a loss of Na^+^ and water balance that contributes to an increased concentration of K^+^ in the hemolymph (hyperkalemia) (MacMillan et al. 2015a; Olsson et al. 2016; Bayley et al. 2018). The combination of cold and hyperkalemia leads to loss of membrane polarization which triggers calcium influx into cells, activating signaling pathways that initiate cell death (MacMillan et al. 2015c; Bayley et al. 2018; Carrington et al. 2020). The increasingly well-resolved physiological basis of chilling injury (see for example (Overgaard et al. 2021)) makes it possible to test both the extent of plastic and evolved changes and also to determine whether any changes in complex phenotypes have similar physiological underpinnings.

*Drosophila melanogaster* are amenable for both field and lab studies and they inhabit a broad range of thermal environments. In the lab, both plastic and evolved increases in basal chill tolerance have been linked to an improved ability to maintain ion homeostasis in the cold. For example, *Drosophila* species that are more chill tolerant and flies that are cold-acclimated both avoid hyperkalemia, in part, through improved renal function in the cold (MacMillan et al. 2015a; Andersen et al. 2017; Yerushalmi et al. 2018). In both cases, these changes to thermal tolerance have been linked to decreased reliance on sodium as an extracellular osmolyte (MacMillan et al. 2015b; Olsson et al. 2016), suggesting that acclimation and evolved differences in chill tolerance are linked to both sodium and potassium regulation. Patterns of evolution in *D. melanogaster* thermal phenotypes have also been well-studied; evolution of *D. melanogaster* populations from high- and low-latitude locations of the east coast of North America demonstrate that populations inhabiting different thermal environments exhibit genetic differences that are likely adaptive in relation to thermal tolerance traits (Mathur and Schmidt 2017). Repeated sampling and common garden rearing of *D. melanogaster* from an orchard in Pennsylvania demonstrated putative adaptive tracking of chill coma recovery from spring to fall, with spring flies showing faster rates of recovery (Behrman et al. 2015).

The extensive research on phenotypic changes of chill coma recovery in *D. melanogaster* makes it possible to combine our laboratory and field approaches to compare the phenotypic effects of acclimation and adaptation across different timescales. To do so, we first assessed the role of acclimation on the phenotypic response to cold exposure using an isofemale line originally derived from London (Ontario, Canada) (Marshall and Sinclair 2010). To compare the magnitude of this variation with that stemming from hundreds of generations of clinal adaptation (Schmidt et al. 2005; Fabian et al. 2012), we collected flies from locations on a well-studied latitudinal gradient in eastern North America (Florida, Pennsylvania, and Maine). Finally, we used a field experiment in which an outbred founder population was introduced into five independent field enclosures to assess adaptive tracking from mid-summer to mid-fall. For each contrast we assessed the fitness-associated phenotype of chill coma recovery time as well as the Na^+^ and K^+^ concentrations that can underlie loss of homeostasis in response to chilling in both males and females. We predicted that the whole-body Na and K content will relate to the cold tolerance (i.e. chill coma recovery time) of *D. melanogaster* in a consistent manner, and that both sexes would consistently differ in ion content as a result of sexual dimorphism. As such, we predict variation in cold tolerance that has 1) been induced within a line via phenotypic plasticity (‘acclimation’), 2) evolved among populations from different climates (‘spatial adaptation’), and 3) evolved rapidly in response to seasonal temperature variation (‘temporal adaptation’) with similar mechanisms of Na and K content associated with improved chill coma recovery across all three contrasts when measured at the organismal level. We further predict that acclimation will have the highest magnitude effects, followed by the spatial adaptation, and adaptive tracking will have the weakest effect.

## Materials and Methods

### Fly lines for the measurement of spatial adaptation and temporal adaptation

Patterns of clinal variation in genotype and phenotype along the east coast of North America are amongst the best studied cases for spatial adaptation in *Drosophila melanogaster* (Schmidt et al. 2008; Telonis-Scott et al. 2011; Bergland et al. 2016). We created five independent outbred populations from Florida, Pennsylvania, and Maine to assess the extent of genetic differentiation in thermal tolerance across latitude. To create each outbred population we randomly selected 10 isofemale lines collected from each latitude and combined 10 males and 10 females from each line into a small cage to allow for recombination. The F1 populations were allowed to expand to >5000 flies and we then collected 100 eggs from each of these populations for the F2 generation. Each population was then held at a minimum population of 300 flies until experimentation. We conducted this protocol 5 times for each latitudinal locality to generate independent outbred populations from each location.

Adaptive tracking in response to seasonal changes within a given locality can also produce pronounced genotypic and phenotypic evolution (Schmidt et al. 2005; Bergland et al. 2014; Behrman et al. 2015). We conducted an experiment in an outdoor orchard facility located at the University of Pennsylvania aimed at assessing the extent of temporal adaptation in response to seasonal change (for more information on the experimental facility see (Rudman et al. 2019)). We released 1000 individuals from an outbred population constructed from 150 isofemale lines originally collected in Pennsylvania into 5 independent 2 m x 2 m x 2 m outdoor mesh enclosures on July 9^th^, 2019 (Grainger et al. 2021). We fed each population 400 mL of modified Bloomington recipe three times weekly. To represent summer adapted flies, we collected ∼5000 eggs from each outdoor cage toward the end of summer (September 12^th^) and kept them reproducing at density-controlled and constant abiotic laboratory conditions until the fall sampling. We collected eggs from each cage again on October 23^rd^ to represent fall adapted fly populations. Temperatures recorded in the field during this time are shown in Figure S1.

### Fly rearing for experiments on thermal acclimation and adaptation

All fly lines were reared in the laboratory in 30 mL glass vials with 7 mL of a banana, corn syrup, agar, and yeast-based medium at 25°C and on a 12 h:12 h L:D cycle. Parental flies from each line were transferred into fresh vials and allowed to lay eggs at 25°C for 24 h, after which the adult flies were removed and the eggs left under the same rearing conditions until adult emergence (at c. 10 days of age), at which point they were moved to vials containing fresh medium. Three days post-emergence, adult flies were sexed under light CO_2_ anesthesia (less than 5 min to avoid effects of CO_2_ on thermal performance; Nilson et al., 2006) and ∼50 flies of each sex were separated and placed in fresh vials.

A separate *D. melanogaster* strain, which was originally collected from London, and Niagara on the Lake Ontario, Canada (Marshall and Sinclair 2010) was also used to investigate long-term acclimation. Flies were maintained in 250 mL plastic bottles containing the same diet and under the same conditions (25°C, 12 h:12 h L:D) as described above. Egg laying and development of experimental flies occurred as described for the other lines. Upon emergence, however, adult flies for the acclimation experiment were split into two groups: half the flies remained at the same conditions (25°C), while the other half were transferred to a 15°C incubator with the same light cycle. Flies matured at their designated acclimation temperature for three days, were sexed (as above), placed in separate vials of ∼50 male and female flies and acclimated for an additional three days. Thus, all flies used for experiments were approximately six to eight days post adult emergence, and acclimated flies spent six days at their respective acclimation temperature.

### Chill coma recovery time

Our chill coma recovery time (CCRT) assay followed previously described methods (MacMillan et al. 2015a). Briefly, flies from each experimental condition (n = 15 flies) were collected and separated individually into 5 mL glass screw-top vials. The vials were sealed, put inside a plastic bag, and then submerged in a mixture of ice and water (0°C) for 6 h. The CCRT was recorded as the time it takes for individual flies to recover a standing position once removed from the ice-water bath and placed at room temperature (22°C) without being disturbed.

### Whole body ion content

Individual flies of each experimental population (n = 15-20 flies) were collected into pre-weighed individual 0.2 mL heat-resistant PCR tubes. Each fly was weighed (inside the tube) using a Sartorius ME5 microbalance (Sartorius Lab Instruments, Goettingen, Germany), and this mass was subtracted from the empty tube mass to determine a wet mass (WM) of the fly. The flies were then dried for ∼24 h in a drying oven at 60°C then reweighed (inside the tube) using the microscale to obtain a dry mass (DM). WM and DM were then used to determine total water content of individual flies.

Whole-body ion contents were measured using flame photometry. First, a stock solution of 100 ppm Li^+^ (Cl was always the counter ion) was prepared from a 3 M Li by dilution with Milli-Q water. Five standard solutions of K^+^ and Na^+^ between 0.1 mM and 5 mM were prepared by dilution with 100 ppm Li, after which 750 µL of each standard was transferred to a conical tube and diluted up to a final volume of 10 mL with 100 ppm Li solution. Standards were prepared in triplicates for measurement in the Sherwood Model 420 Flame Photometer (Sherwood Scientific, Cambridge, UK). After calibration with a blank solution of 100 ppm Li, K^+^ and Na^+^ concentrations were recorded from five prepared standards to create K^+^ and Na^+^ standard curves. Fly samples were prepared by transferring the dried flies to 1.7 mL microcentrifuge tubes and grinding each fly using a mortar and pestle before adding 200 µL of 100 ppm Li to the tube. The samples were homogenized using a sonicator (Qsonica, Newton, CT, USA), until the solution was visibly homogenous. The tubes were then centrifuged at 10,000 × *g* for 5 min at 4°C. After centrifugation, 150 µL of the supernatant was transferred to a 2 mL microcentrifuge tube and diluted up to a final volume of 2 mL with 100 ppm Li solution. The pellet was discarded. Na^+^ and K^+^ concentrations in the diluted supernatants were measured using the flame photometer and converted to [Na^+^] and [K^+^] in each fly (interpreted as an averaging of all concentrations in all parts of the fly) by reference to the standard curves and accounting for initial water content. Na and K content were also expressed relative to dry mass (rather than water content) to elucidate whether ion content or water content was driving differences in mean concentration. This was done by converting concentrations to molar quantities of each element and accounting for the individual fly’s dry mass.

### Data analysis

All data analysis was done in the R language for statistical computing (version 3.0.1) (R Development Core Team 2019). We tested for differences in chill coma recovery time (CCRT) among acclimated, climate-adapted or seasonally-adapted *D. melanogaster* using mixed-effects models using the lme() function (Pinheiro et al. 2020) with treatment/climate/season, and sex treated as fixed effects and with line and sampling date treated as random effects. Similarly, we tested for differences in ion concentration (in mM) and ion content (in mmol/mg) of sodium and potassium obtained by flame photometry using mixed-effects models with the same model form. We also tested for differences in water content (as a proportion of body mass) using measurements of wet and dry mass, and analyzed these differences using the same mixed-effects models. In all cases, sampling date had no effect on our results, so we simplified our analysis by removing collection date as a random effect. We found intriguing patterns in the relationship between ion and water content in acclimated flies and tested whether ion content was tied to variation in water content (as a proportion of body mass) separately in male and female acclimated files using generalized linear models with water content and acclimation temperature treated as factors. To test for the same patterns in climate and seasonally adapted flies we used mixed effects models with treatment/climate/season, and sex treated as factors and with line treated as a random effect.

## Results

### Chill coma recovery time

We quantified cold tolerance in all lines and both acclimation groups using chill coma recovery time (CCRT). Across all three groups (acclimation, spatial-, and temporal adaptation), we noted a clear pattern of females recovering from chill coma 5-15 minutes faster than males within the same line/acclimation group (Figure 1). Cold acclimation reduced CCRT by approximately 19 min in males and 16 min in females, climate-adaptation reduced CCRT by approximately 9.5 min in males and 5 min in females (FL vs. ME), and seasonal adaptation altered CCRT by less than 1 min in both males and females.

**Figure 1.**
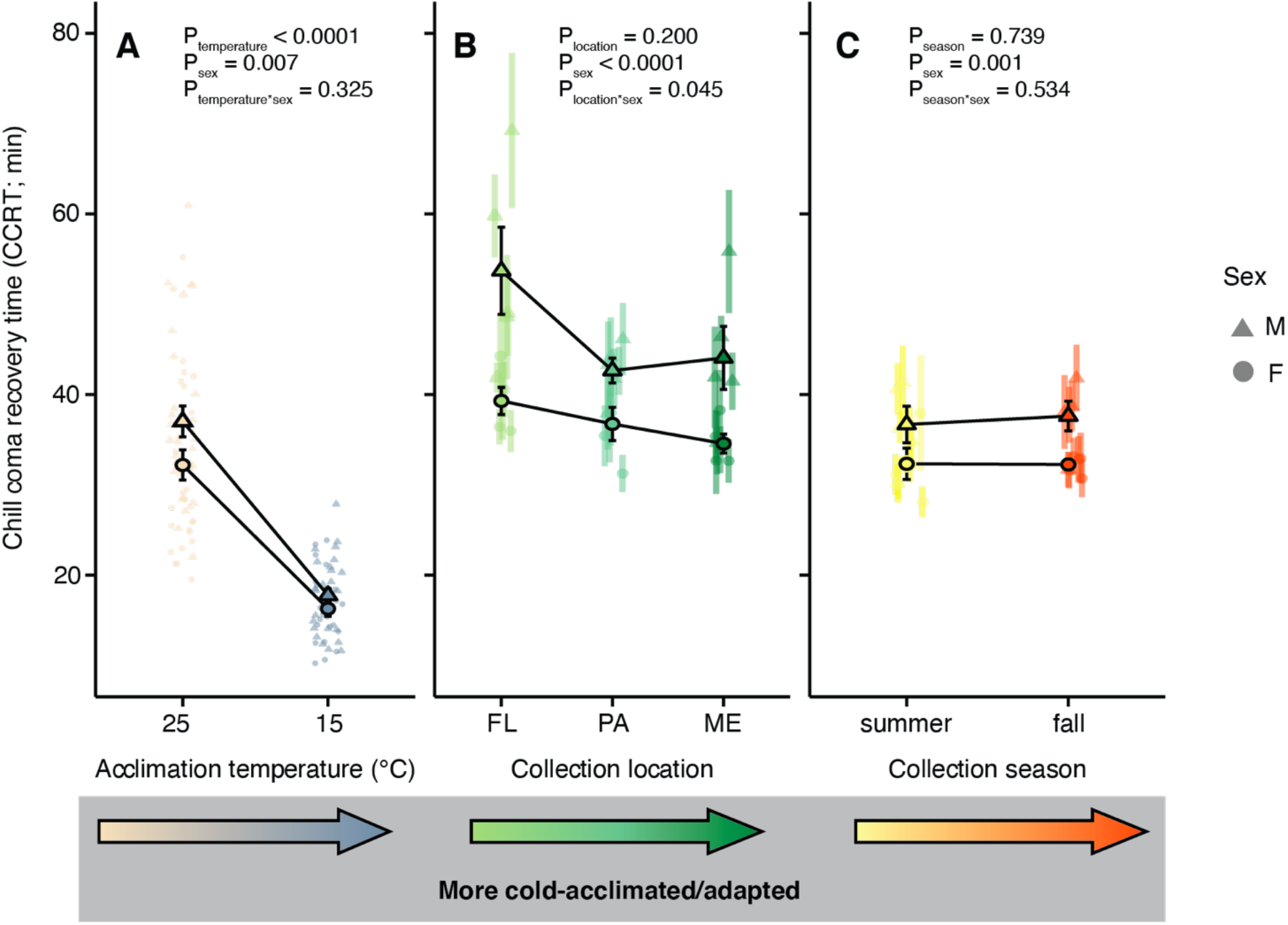
Acclimation and collection location, but not season relate to cold tolerance quantified as chill coma recovery. CCRT of male (triangles) and female (circles) *D. melanogaster* following exposure to 0°C for 6 h. Flies have undergone acclimation (A) or were collected across latitudes that vary considerably in temperature (B) or in the season of collection (C). Opaque symbols with error bars represent global means (± sem) among individuals (A) or lines (B and C). Error bars that are not visible are obscured by the symbols. P-values calculated using mixed-effects models in R (see Table 1 for statistics).

In the acclimated flies, there was a significant main effect of cold-acclimation on CCRT (*P* < 0.0001), as well as a main effect of sex (*P* = 0.008; Table 1; Figure 1A).

**Table 1.** Summary of results from mixed-effects models on chill coma recovery time (CCRT). Effects of acclimation temperature, climate of collection, or season of collection and sex are shown for lines of *D. melanogaster* that have been acclimated, spatially adapted or temporally adapted (respectively). For flies collected from different climates or in different seasons, mixed-effects models included replicate line as a random effect. Shaded rows indicate significance based on *P*<0.05.

| Group | Effect | F | P-value | df |
| --- | --- | --- | --- | --- |
| Acclimation | Acclimation Temp. | 178.52 | < 0.0001 | 1, 106 |
| Acclimation | Sex | 7.38 | 0.008 | 1, 106 |
| Acclimation | Acclimation Temp. * Sex | 0.98 | 0.325 | 1, 106 |
| Spatial | Climate | 1.85 | 0.200 | 2, 12 |
| Spatial | Sex | 45.21 | < 0.0001 | 1, 501 |
| Spatial | Climate * Sex | 3.12 | 0.045 | 2, 501 |
| Temporal | Season | 0.12 | 0.739 | 1, 8 |
| Temporal | Sex | 11.42 | 0.001 | 1, 340 |
| Temporal | Season * Sex | 0.39 | 0.534 | 1, 340 |

In the spatial adaptation flies, the CCRT was significantly affected by sex (*P* < 0.0001), and there was no significant main effect of origin location (*P* = 0.200). There was, however, a significant interaction in the effects of location and sex on CCRT (P = 0.045; Table 1; Figure 1B). Female flies from the more poleward climate (Maine; ME) recovered ∼ 5 min faster from chill coma than Florida (FL) females reared under the same conditions. By contrast, in male spatial adaptation fly lines, the Pennsylvania (PA) lines had the fastest CCRT while the FL males recovered slowest (a difference of ∼ 12 min).

In the temporal adaptation flies (summer vs. fall), there was no significant effect of adaptive tracking across season (*P* = 0.739), but there was a significant effect of sex on CCRT (*P* = 0.001; Table 1; Figure 1C).

### Ion and water content

We quantified Na and K content in *D. melanogaster* (expressed per mg of dry mass) using flame photometry on individual flies that were weighed before and after drying to determine water content. Neither thermal acclimation (*P* = 0.922) nor sex (p = 0.517) influenced Na content in acclimated flies (Table 2; Figure 2A). By contrast, K content was significantly higher in cold acclimated flies (*P* < 0.0001). There was no main effect of sex (*P* = 0.482), nor was there an interactive effect of acclimation temperature and sex on K content (P = 0.468; Table 2; Figure 2D). K content increased by ∼ 0.1 μmol mg^-1^ (∼ 19 % increase) in both the male and female flies that acclimated at 15°C compared to the male and female flies that acclimated at 25°C. There were no significant effects of acclimation temperature (p = 0.148) or sex (p = 0.122) on water content in the acclimated flies (Table 2; Figure 2G).

**Table 2.** Results of mixed effects models of ion and water content (WC) in acclimated, climate-adapted, or seasonally-adapted fly lines. Shaded rows indicate significance based on *P*<0.05.

| Variable | Group | Effect | F | P-value | df |
| --- | --- | --- | --- | --- | --- |
| Na | Acclimation | Temperature | 0.01 | 0.922 | 1, 94 |
| Na | Acclimation | Sex | 0.42 | 0.517 | 1, 94 |
| Na | Acclimation | Temperature * Sex | 1.65 | 0.203 | 1, 94 |
| Na | Spatial | Climate | 0.03 | 0.969 | 2, 12 |
| Na | Spatial | Sex | 6.59 | 0.011 | 1, 408 |
| Na | Spatial | Climate * Sex | 13.85 | <0.0001 | 2, 408 |
| Na | Temporal | Season | <0.01 | 0.980 | 1, 8 |
| Na | Temporal | Sex | 17.21 | <0.0001 | 1, 276 |
| Na | Temporal | Season * Sex | <0.01 | 0.999 | 1, 276 |
| K | Acclimation | Temperature | 34.63 | <0.0001 | 1, 94 |
| K | Acclimation | Sex | 0.50 | 0.482 | 1, 94 |
| K | Acclimation | Temperature * Sex | 0.53 | 0.468 | 1, 94 |
| K | Spatial | Climate | 1.15 | 0.348 | 2, 12 |
| K | Spatial | Sex | 2.02 | 0.156 | 1, 408 |
| K | Spatial | Climate * Sex | 18.14 | < 0.0001 | 2, 408 |
| K | Temporal | Season | 0.08 | 0.780 | 1, 8 |
| K | Temporal | Sex | 2.40 | 0.123 | 1, 276 |
| K | Temporal | Season * Sex | 6.78 | 0.010 | 1, 276 |
| WC | Acclimation | Temperature | 2.13 | 0.148 | 1, 94 |
| WC | Acclimation | Sex | 2.44 | 0.122 | 1, 94 |
| WC | Acclimation | Temperature * Sex | 0.07 | 0.790 | 1, 94 |
| WC | Spatial | Climate | 1.99 | 0.180 | 2, 12 |
| WC | Spatial | Sex | 12.86 | 0.0004 | 1, 408 |
| WC | Spatial | Climate * Sex | 24.00 | <0.0001 | 2, 408 |
| WC | Temporal | Season | <0.001 | 0.991 | 1, 8 |
| WC | Temporal | Sex | 34.17 | <0.0001 | 1, 276 |
| WC | Temporal | Season * Sex | 0.01 | 0.922 | 1, 276 |

**Figure 2.**
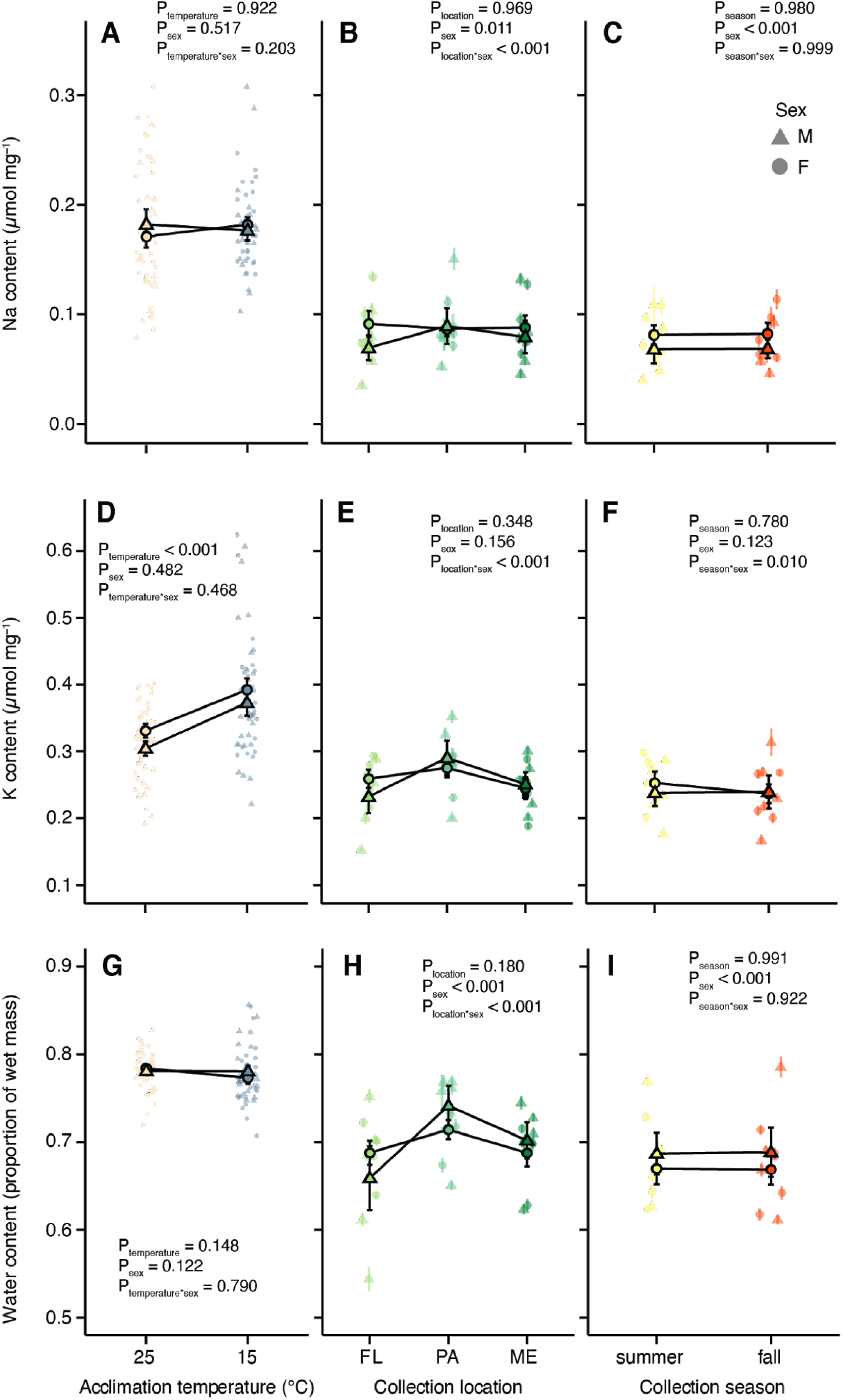
Acclimation alters whole body K content without changing water content, while climate adaptation is associated with sex-specific differences in both K and water storage. Na and K content of male (triangles) and female (circles) obtained by flame photometry and converted to values of ion content (in mmol) per mg of dry mass using standard curves of Na and K from 0.1 – 5 mM, while water content was calculated as a proportion of wet mass. Flies have undergone acclimation (A; D; G) or are cold adapted based on collection location (B; E; H) or season of collection (C; F; I). Opaque symbols with error bars represent global means (± sem) among individuals (A; D; G) or lines (B; E; H and C; F; I). Error bars that are not visible are obscured by the symbols. P-values were calculated using mixed-effects models in R.

In the spatially adapted flies, there was a significant interactive effect of sex and origin on Na content (*P* < 0.0001; Table 2; Figure 2B). Flies from Pennsylvania (PA) had a higher sodium content than flies from Maine (ME) or Florida (FL). There was also a significant main effect of sex (*P* = 0.011), however unlike in the cold acclimated flies, spatially adapted females tended to have a lower Na content across all climates. In these flies, K content displayed a similar trend to Na, where flies from PA contained more K and those from FL contained less (by ∼ 0.05 μmol mg^-_1_^). There was a significant interactive effect of origin location and sex on K content (*P* < 0.0001; Table 2; Figure 2E), but neither location (*P* = 0.348) nor sex (*P* = 0.156) had a main effect on K content. The water content of flies collected from different locations was significantly affected by an interaction between location and sex (*P* < 0.0001; Table 2; Figure 2H); males from PA and FL carried more water than females, but males from ME carried less water than females from the same location. Lines collected in Florida tended to carry less water than those from the more poleward climates, and both male and female flies from Pennsylvania carried the most water.

In temporal adaptation fly lines, we found a significant effect of sex (*P* < 0.0001), but not collection season (*P* = 0.980) on Na content (Table 2; Figure 2C). Female flies had higher Na content across both seasons when compared to the male flies. Season and sex interacted to influence K content (*P* = 0.010); females had greater K content in the summer but this difference was absent from flies collected in the fall (Figure 2F). There were no main effects of either season (*P* = 0.780) nor sex (*P* = 0.123) on K content (Table 2; Figure 2F). The water content of temporal adaptation flies differed only according to sex (*P* < 0.0001; Table 2; Figure 2I), with males holding more water than females regardless of the season during which the lines were sampled.

### Relationship between ion content and cold tolerance

Changes in ion and water content can both influence physiologically-relevant ion concentrations, and Na and K are almost exclusively retained in animals in their free (unbound) ionic forms (Na^+^, K^+^). In Figure 3, we examined the relationships between cold tolerance (CCRT) and whole-body average [K^+^] ([Na^+^] in Figure S2). While improvements in cold tolerance with cold acclimation were associated with an increase in [K^+^], the opposite was true in spatial adaptation flies (Figure 3), and temporal adaptation flies had no distinct relationship between the potassium concentration and CCRT (but also very little difference in CCRT). These relationships also appear to differ somewhat according to sex (Figure 3).

**Figure 3.**
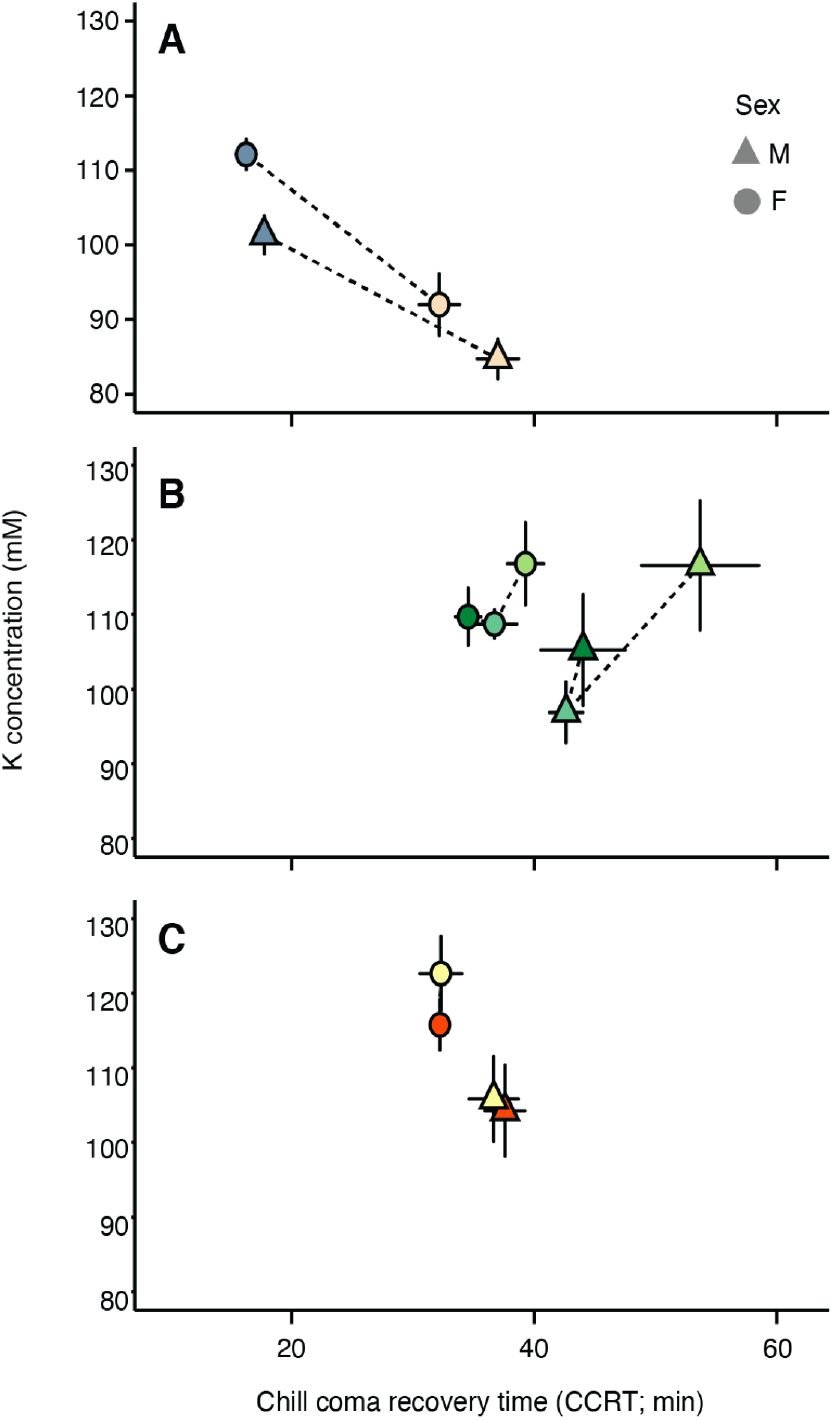
Cold acclimation was associated with increased whole body [K^+^], while flies from colder climates tended to have lower, rather than higher, average [K^+^], because of the influence of water. Relationships between chill coma recovery time and average whole-body potassium concentration in both male (triangles) and female (circles) *D. melanogaster*. Means (± sem) among individuals following acclimation (A) or among lines collected at different locations (B) or collected in different seasons (C) are shown.

The relationship between [Na^+^] and cold tolerance displayed relatively similar trends to the potassium concentration (Figure S2). The acclimated flies overall tended to have a negative relationship between the [Na^+^] and cold tolerance, but there was only a ∼ 5 mM difference in concentration between the warm- and cold-acclimated flies (Figure S2). As with the potassium/cold tolerance relationship, the spatial adaptation flies displayed a positive trend between the sodium concentration and CCRT, while the temporal adaptation flies had no obvious relationship between the sodium concentration and cold tolerance (Figure S2).

### Relationships between ion and water content

Both Na^+^ and K^+^ are critically important to water balance as well as cold tolerance. We noted that relationships between these ions and water content differed among experimental groups, and this was most striking in regard to acclimation and in female flies. Warm acclimated flies preferentially retained Na^+^ along with water; there was a positive relationship between water content and Na content in both males and females (Figure 4A) but not K content (Figure 4B). However, in the cold acclimated flies, the opposite relationship is clear; sodium content and water content had no distinct relationship (Table S1), whereas potassium content increased along with water content (Figure 4). This discrepancy is clear from significant interactions in the effects of acclimation temperature and water content on ion content (P<0.006) in all cases except for Na in males where it was near-significant (P = 0.062) and following the same trend. To remove any possible influence of body size variation, we calculated the ratio of Na:K and found that specifically in females, cold acclimated flies retain more Na relative to K as whole-body water content increases, while warm acclimated flies do the opposite (P<0.001; Figure 4E). While the trend was similar, there was no significant effect of water content on the Na:K ratio in male acclimated flies (P = 0.107).

**Figure 4.**
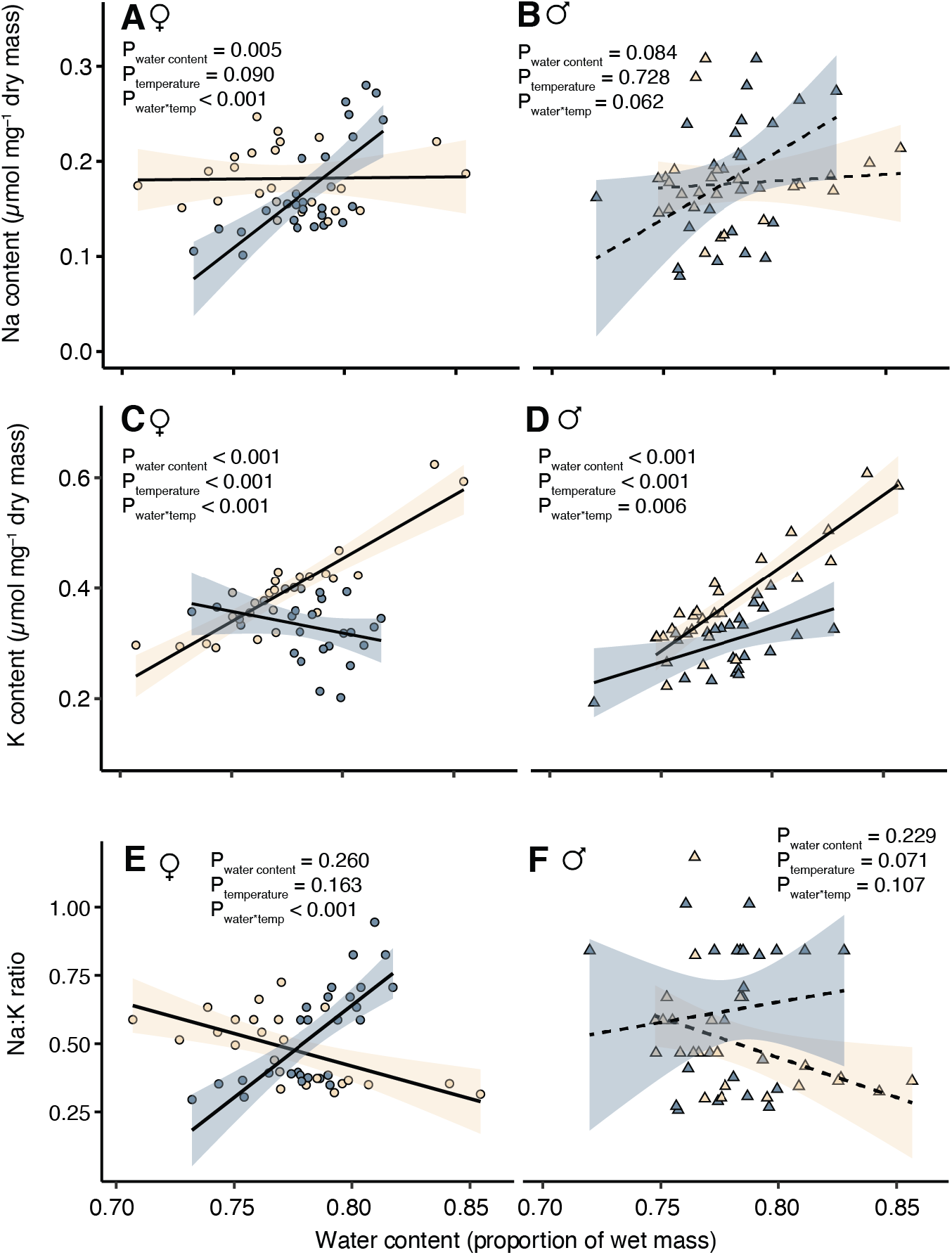
Relationships between water content and ion content in warm (yellow) and cold (blue) acclimated female (A,C; circles) and male (B,D; triangles) flies. In both males and females, higher water content was generally associated with accumulation of K in warm acclimated flies, but Na in cold-acclimated flies (interaction effects). Solid black lines (dashed if interaction term is not significant) with transparent shaded areas represent linear models to demonstrate trends (mean and sem of parameter estimates). Full results of generalized linear models are presented in Table S1.

Notably, in both the temporal and spatial adaptation fly lines, we found no such preference for one ion over the other in flies that had higher water content; both ions increased in abundance in flies that carried more water and there was no significant effect of origin location or season in either sex (Figure S3-S4; Table S2). In both temporal and spatial adaptation lines, K content was more tightly associated with water content than Na (Table S2).

## Discussion

### Acclimation and adaptation of basal cold tolerance shape whole animal ion content differently

Here, we sought to compare the phenotypic and physiological differences among flies that experienced long-term thermal acclimation, and further compare differences in the same traits to evolved variation in populations originating from different latitudes and seasons. We used a common chill tolerance assay (CCRT; Figure 1), and measured whole body Na and K, two ions which play key roles in the cold tolerance of *Drosophila* (Figure 2) (MacMillan and Sinclair 2011; MacMillan et al. 2015b; Overgaard and MacMillan 2017). Long-term acclimation at 15°C decreased CCRT to a greater degree than the observed variation in CCRT among lines of flies adapted to different climates (Figure 1). We also saw little evidence for variation in CCRT arising as a result of temporal adaptation (Figure 1). Variation in chill tolerance is well studied in *Drosophila* in relation to both acclimation (Sinclair and Roberts 2005; Overgaard et al. 2008; Koštál et al. 2011; Findsen et al. 2013) and climatic and seasonal variation (Gibert et al. 2001; Hoffmann et al. 2002; Overgaard et al. 2011; Kellermann et al. 2012; MacMillan et al. 2016; MacLean et al. 2019) and the variation observed is a prerequisite for asking questions about how ion balance and chill tolerance co-vary.

As is the case with CCRT, our warm- and cold-acclimated acclimated flies also displayed the largest differences in total ion content when compared to populations originating from different latitudes or seasons. This was most evident from whole-body potassium content, which increased in cold-acclimated flies (Figure 2D) while they maintained the same water content as the warm-acclimated flies (Figure 2G). This (more K in the same amount of water) drove an overall increase in whole body average [K^+^] in both male and female cold-acclimated flies. Interestingly, this pattern is wholly different from how changes in ion balance matched with the chill tolerance of flies adapted to different climates; in this group, whole body K content was largely stable regardless of origin location (Figure 2E), while water content varied (Figure 2H). This pattern resulted in flies from cooler climates having overall lower, rather than higher, whole body [K^+^] (the same amount of ions in a larger water volume; Figure 3B). Taken together, acclimation and latitudinal adaptation of basal tolerance appear to be related to whole animal average [K^+^] in opposing directions: while improved cold tolerance is associated with higher [K^+^] in acclimated flies, it is instead associated with lower [K^+^] in cold climate adapted flies.

We note that here, sex had detectable effects on both CCRT and ion balance. Research on *Drosophila* cold tolerance has been mainly focused on females, but male and female flies respond to chilling differently (David et al. 1998; Vesala and Hoikkala 2011; Rajpurohit and Schmidt 2016). As observed here, females are generally more cold tolerant, as they recovered from chill coma faster in all groups (Figure 1), and this difference may reflect sex-specific differences in expression of reproductive dormancy (Kurogi et al. 2021). Females also varied among groups in the relationships between ion content and water content, but we specifically note that in all three groups (acclimation, clinal, and seasonal adaptation), females always tended to have higher whole-body average [K^+^] relative to males, meaning females carried more K per unit body water (Figure 3). This suggests that sex effects on K^+^ balance relate to cold tolerance in the same manner as acclimation (higher [K^+^] is associated with greater cold tolerance), but differently from adaptation of basal tolerance.

A wide-ranging literature documenting rapid adaptation has shaped thinking about the importance and pace of adaptation in response to environmental change (Sanderson et al. 2022). This includes a seasonal adaptation experiment from which the temporal adaptation flies studied here were taken, which found rapid phenotypic adaptation in response to both seasonal change and the presence of a competitive species (Grainger et al. 2021). A growing number of experiments tracking *Drosophila* populations through time have found evidence of rapid adaptation (Rajpurohit et al. 2018, Rudman et al. 2019, Rudman et al. 2022). Moreover, patterns of spatial intraspecific genetic variation across *Drosophila* populations are well-documented and the resulting phenotypic variation between populations can be considerable (Hoffman et al. 2002, Schmidt et al. 2005, Fabian et al. 2012). Here we find modest evidence for spatial adaptation in cold tolerance phenotypes and no evidence for rapid temporal adaptation in response to a fall acclimatization. There are several putative explanations for the lack of cold adaptation in seasonally cooling conditions – chief among them that the temperature drop prior to sampling had been modest relative to what is observed later in the fall (Fig. S2). Indeed, a recent paper in which rapid evolution of chill coma recovery was observed across seasonal time did not find significant evolution over the window sampled for this study (Rudman et al. 2022). Perhaps cooling temperatures was not a strong enough agent of selection over a long enough duration to drive adaptation during the time period studied here. More broadly, it is also possible that high-profile cases of rapid adaptation are creating an expectation of ubiquitous adaptation: A recent meta-analysis of experiments found no evidence for consistent life history trait adaptation in response to manipulative climatic warming (Grainger and Levine 2021). Our data support a view in which acclimation is a singular mode by which phenotypes shift in response to rapid environmental change. Future synthetic work comparing the predictability and magnitude of plastic and evolved changes will be key in identifying the mechanisms by which organisms are most likely to respond to abiotic challenges.

The observation that thermal plasticity and adaptation of basal thermal tolerance are associated with changes in ion stoichiometry may point us toward important and undescribed mechanisms of thermal performance in insects. Ultimately, however, connecting the observed changes in K and water content to current knowledge on the physiological mechanisms of cold tolerance is challenging when the changes are observed at the whole organism level. One of the most pressing questions that emerges from this work, therefore, is what organs in the body are driving the observed variation in stoichiometry. In most insects, including *Drosophila*, Na^+^ is the primary extracellular ion and an important osmolyte, and intracellular [Na^+^] levels are lower in the cell cytoplasm (∼18 mM) than the hemolymph (∼70 mM; MacMillan et al. 2015b). By contrast, [K^+^] is higher inside cells (∼160 mM) and maintained at low levels (∼10-15 mM) in the hemolymph. When chilled, chill-susceptible insects like *D. melanogaster* lose the ability to maintain these gradients and ion balance is progressively lost as Na^+^ leaks across epithelia (osmotically taking water with it) and [K^+^] concurrently rises in the hemolymph (Koštál et al. 2006; MacMillan and Sinclair 2011; MacMillan et al. 2014; Andersen and Overgaard 2019). If severe enough, high [K^+^] in the hemolymph depolarizes cells, causing catastrophic Ca^2+^ influx and cell death (ultimately causing organismal death; (MacMillan et al. 2015c; Bayley et al. 2020; Carrington et al. 2020)). Similarly, a rapid loss of ion balance occurs within the central nervous system when insects like *D. melanogaster* are sufficiently cooled, albeit it *via* an unknown, different physiological mechanism with unknown consequences for survival and fitness (Robertson et al. 2017, 2020). Because a loss of systemic ion balance is an important driver of low temperature injury and death (Overgaard and MacMillan 2017), variation in chilling tolerance within and among species is thought to be closely tied to the magnitude of ion (most notably Na^+^ and K^+^) gradients across membranes and epithelia *before* a stress is applied, as well as how cells, tissues, and organs responsible for maintaining ion gradients at the local and organismal level respond *during* chilling (MacMillan et al. 2015a, 2017; Yerushalmi et al. 2018; Andersen and Overgaard 2020). Specifically, the status of [Na^+^] gradients before chilling occurs appears relevant to both cold acclimation and cold adaptation among species; cold acclimated *D. melanogaster* and cold adapted *Drosophila* species maintain lower [Na^+^] gradients without altering hemolymph osmolarity (MacMillan et al. 2015b; Olsson et al. 2016), such that water balance may be better maintained in the cold. For this reason, we expected to see adaptive changes in whole-body Na content in the present study, but our results instead suggest that changes in these gradients may occur without any gain or loss of organismal ion content. Total body Na content may nonetheless intersect in important ways in insects to influence thermal performance and other fitness-related traits in insects; dietary Na^+^ can alter insect growth and may be limiting in some food webs (Welti et al. 2019; Peterson et al. 2021).

Given that a loss of K^+^ balance is a direct cause of chilling injury, we did not expect cold acclimated flies to retain a greater amount of K than their warm-acclimated counterparts (*sensu* Figure 3). Whether or how this additional K is contributing to improved chill tolerance by altering K^+^ gradients entirely depends on the location in which it is being accumulated. Increased intracellular [K^+^] without any change in membrane ion permeability or extracellular [K^+^] would cause hyperpolarization of nerve and muscle cells, which may protect against the cold-induced depolarization responsible for chilling injury. Establishing and maintaining this state would likely come at an energetic cost (and many secondary consequences) that might not be favourable to maintain constitutively when conditions are warmer (as in the case of cold-adapted fly lines maintained at constant 25°C). By contrast, additional K^+^ in the lumen of the digestive tract could establish gradients that make K^+^ clearance from the hemolymph *via* the gut epithelia and renal system less energetically favourable, which would be detrimental to chill tolerance. Similarly, storing additional K^+^ in the hemolymph seems maladaptive (as the goal is to prevent hemolymph hyperkalemia), but recent findings suggest that the depolarizing effects of a putative cold-acclimation-induced increase in hemolymph K^+^ concentration could be mitigated by an increased expression of K^+^ channels in the cell membranes (Bayley et al. 2020). Lastly, there is the possibility that the increase in whole-fly K^+^ concentration after cold acclimation is the result of a wholesale increase in K^+^ content in all bodily compartments, which, in the light of the current conceptual model for insect chill susceptibility (Overgaard and MacMillan 2017), would be detrimental for the maintenance of membrane potential, and by extension survival, unless counterbalanced by a suite of other (potentially costly) adaptive responses (e.g. differential expression of channels and/or ion transporters). Importantly, changes in ion content at the organismal level may arise not just from changes in ion concentrations, but also from changes in the relative size of organs and extracellular fluids within the body. In this scenario, cold-acclimated or cold adapted flies may retain more or less total K (respectively) because of underlying variation in the relative investment in organs that contain more or less K per unit mass.

### Conclusion

Overall, the findings presented here arise from collaborative field and lab approaches and perspectives from both comparative evolutionary genomics and integrative and comparative physiology. We found that phenotypic plasticity and spatial adaptation can both shape cold tolerance, but that acclimation produced the greatest magnitude change in this trait in *D. melanogaster*. Given that ion balance is critical to chill tolerance in insects, changes to ion uptake and elimination processes may drive the differences in whole organism ion stoichiometry that we observed here, and thereby alter thermal performance. While we have described these changes and how they can differ at the whole-animal level, there remains a significant gap in our understanding of how these changes relate to ion gradients across cell membranes and epithelia, and how, precisely, such changes modulate low temperature performance traits. In the present study we treated acclimation and special and temporal variation in basal tolerance as separate, but it remains unclear how the capacity for plasticity can itself evolve in response to change, and to what degree the same physiological mechanisms underlie differences in acclimation capacity and basal tolerance.

## Supporting information

Data archive

## Data Availability

All data is provided as a supplementary file for review and the same file will be included as supplementary material or uploaded to Dryad should the manuscript be accepted for publication.

---

**Figure S1:**
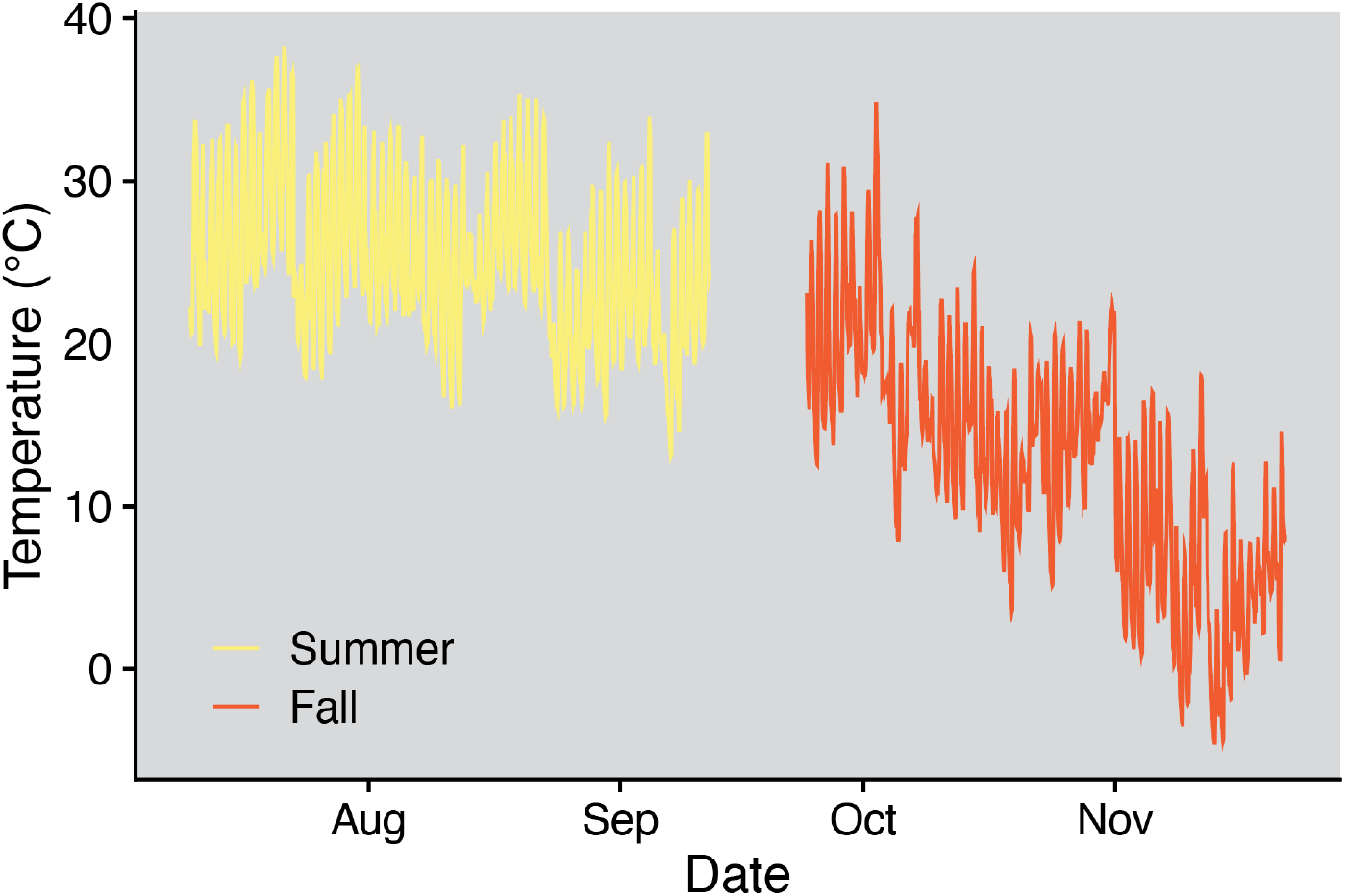
Average hourly temperature from the seasonal adaptation experiment as measured by 8 independent temperature loggers.

**Figure S2.**
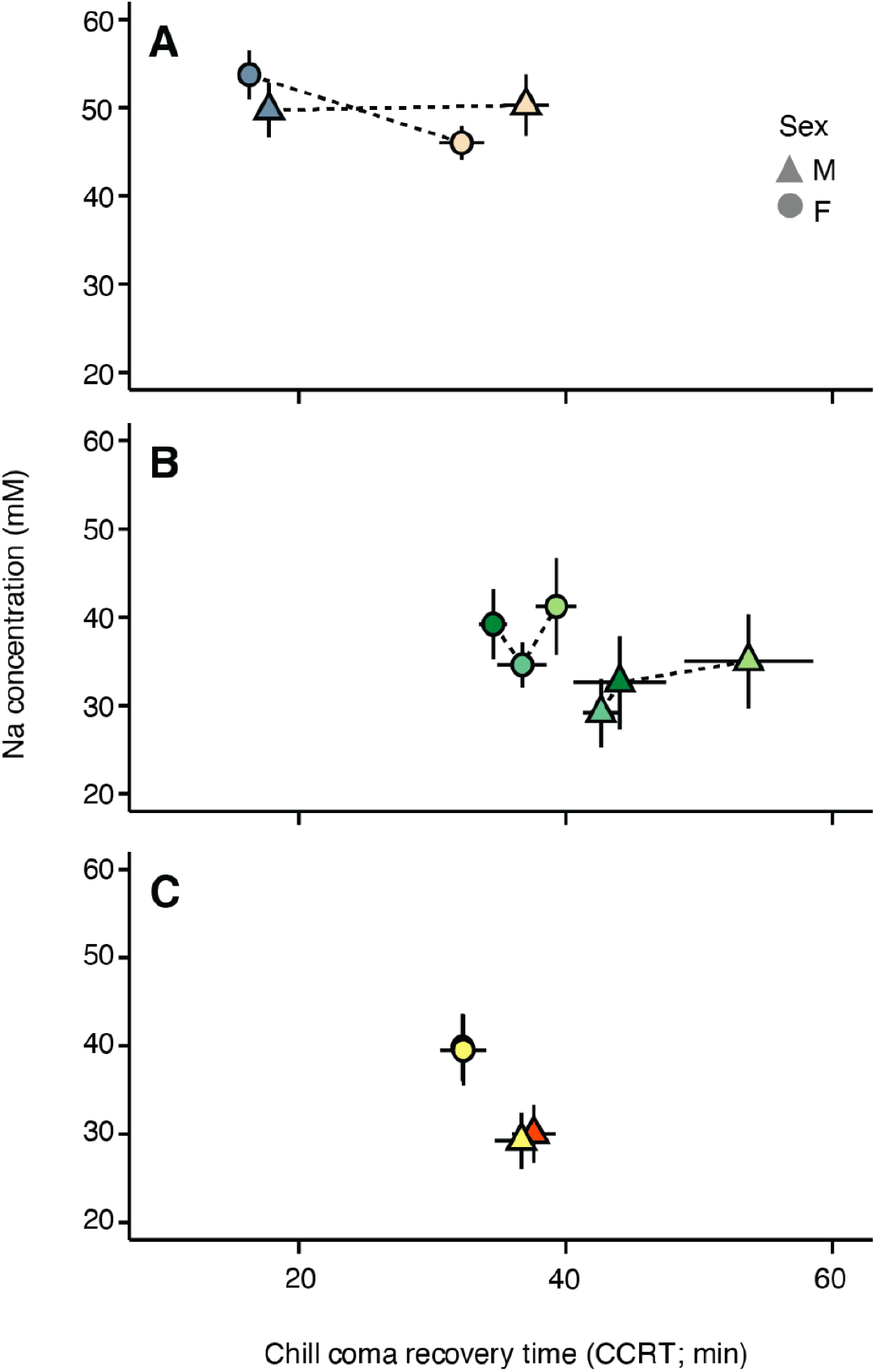
Relationships between chill coma recovery time and average whole-body sodium concentration in both male (triangles) and female (circles) *D. melanogaster*. Means (± sem) among individuals following acclimation (A) or among lines collected at different locations (B) or collected in different seasons (C) are shown.

**Figure S3.**
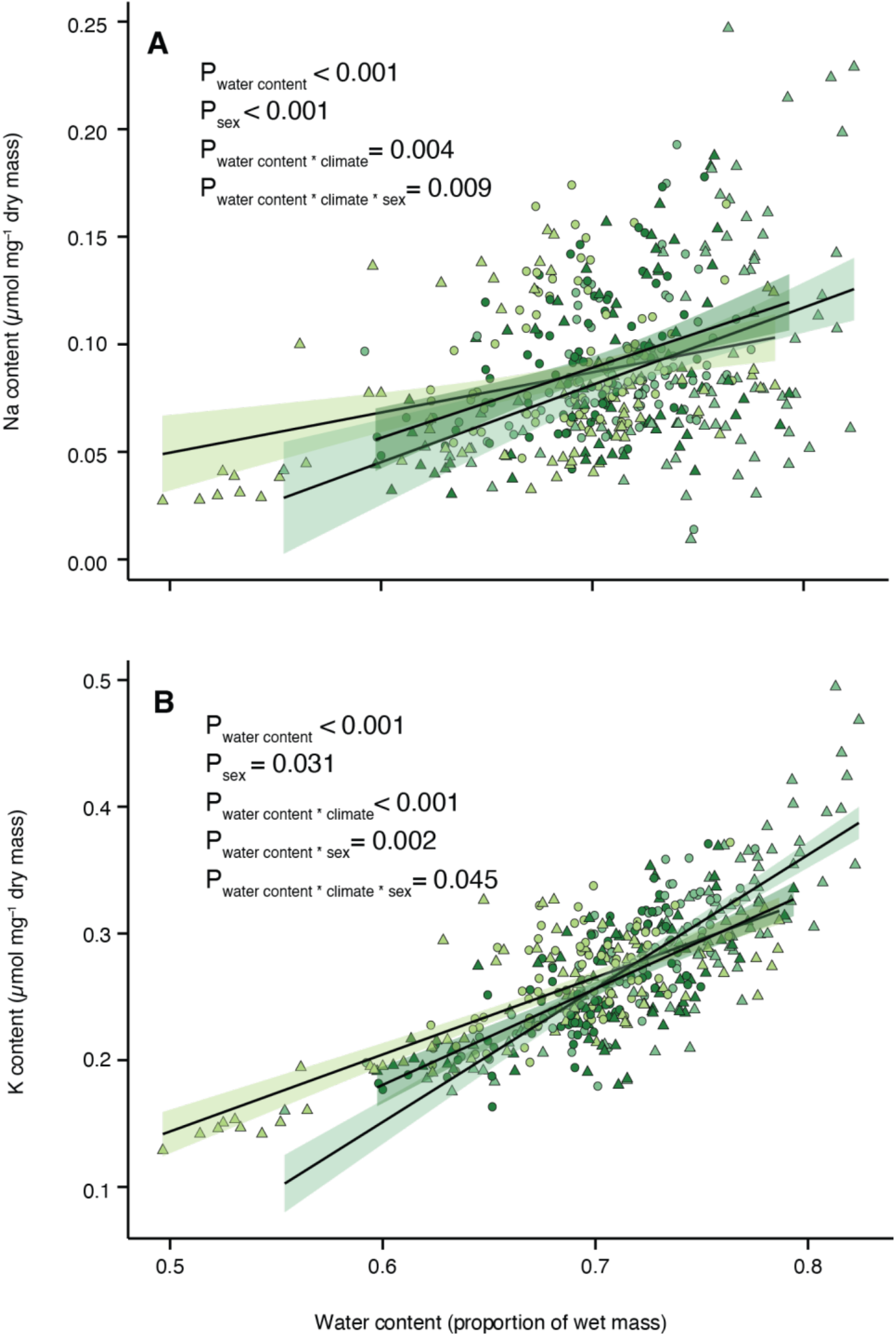
Relationship between sodium or potassium content and water content in male (triangles) and female (circles) climate-adapted *D. melanogaster*. Solid black lines with transparent shaded area represent linear models to demonstrate trends (mean and sem of parameter estimates). Full results of mixed effects models are presented in Table S2.

**Figure S4.**
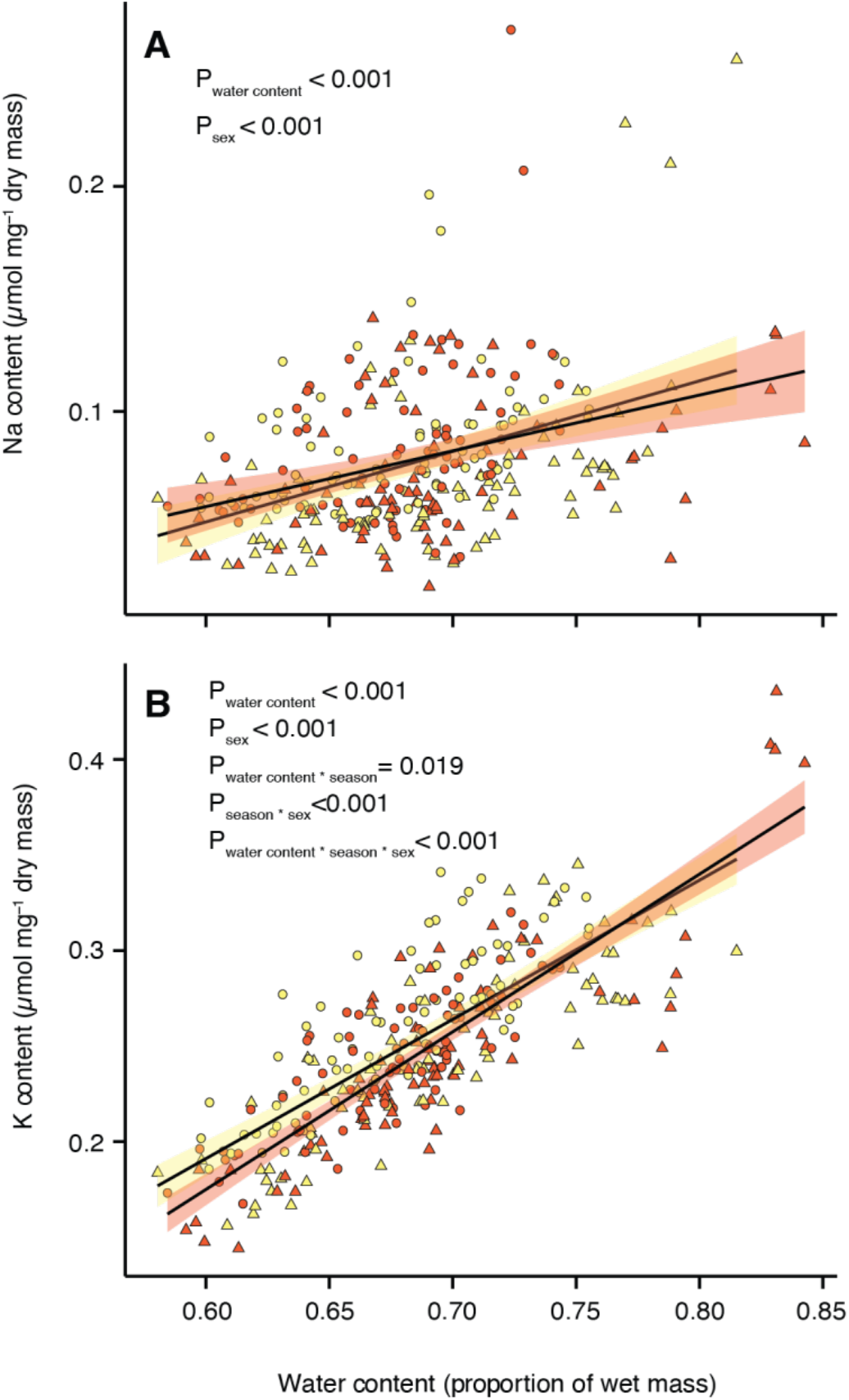
Relationship between sodium or potassium content and water content in male (triangles) and female (circles) seasonally-adapted *D. melanogaster*. Solid black lines with transparent shaded area represent the overall data trend.

**Table S1.**
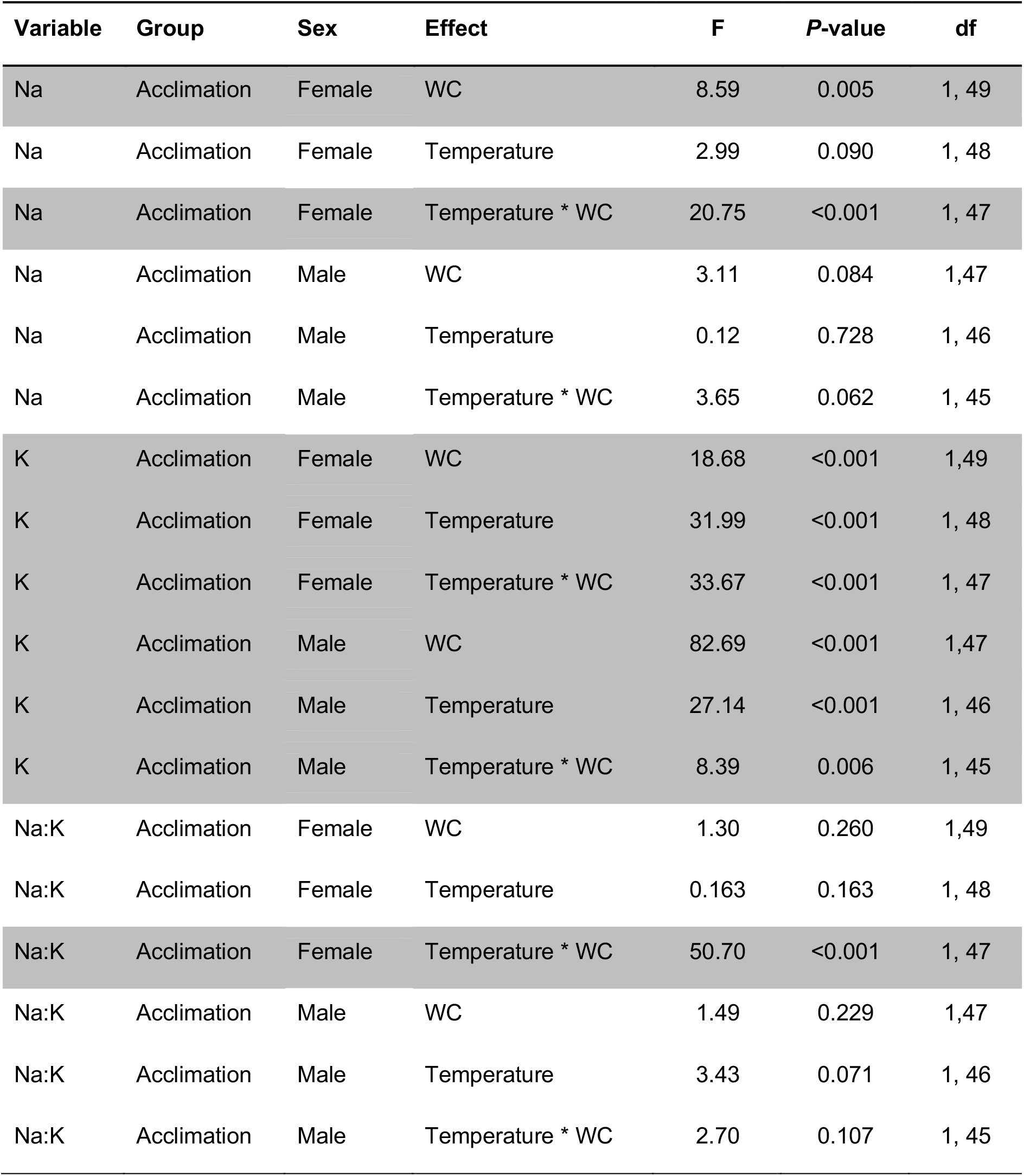
Results of generalized linear models and mixed effects models of ion content relative to water content (WC) in acclimated, climate-adapted, or seasonally-adapted fly lines. Shaded rows indicate significance based on *P*<0.05. Na:K = Ratio of Na relative to K in body.

| Variable | Group | Sex | Effect | F | P-value | df |
| --- | --- | --- | --- | --- | --- | --- |
| Na | Acclimation | Female | WC | 8.59 | 0.005 | 1, 49 |
| Na | Acclimation | Female | Temperature | 2.99 | 0.090 | 1, 48 |
| Na | Acclimation | Female | Temperature * WC | 20.75 | <0.001 | 1, 47 |
| Na | Acclimation | Male | WC | 3.11 | 0.084 | 1, 47 |
| Na | Acclimation | Male | Temperature | 0.12 | 0.728 | 1, 46 |
| Na | Acclimation | Male | Temperature * WC | 3.65 | 0.062 | 1, 45 |
| K | Acclimation | Female | WC | 18.68 | <0.001 | 1, 49 |
| K | Acclimation | Female | Temperature | 31.99 | <0.001 | 1, 48 |
| K | Acclimation | Female | Temperature * WC | 33.67 | <0.001 | 1, 47 |
| K | Acclimation | Male | WC | 82.69 | <0.001 | 1, 47 |
| K | Acclimation | Male | Temperature | 27.14 | <0.001 | 1, 46 |
| K | Acclimation | Male | Temperature * WC | 8.39 | 0.006 | 1, 45 |
| Na:K | Acclimation | Female | WC | 1.30 | 0.260 | 1, 49 |
| Na:K | Acclimation | Female | Temperature | 0.163 | 0.163 | 1, 48 |
| Na:K | Acclimation | Female | Temperature * WC | 50.70 | <0.001 | 1, 47 |
| Na:K | Acclimation | Male | WC | 1.49 | 0.229 | 1, 47 |
| Na:K | Acclimation | Male | Temperature | 3.43 | 0.071 | 1, 46 |
| Na:K | Acclimation | Male | Temperature * WC | 2.70 | 0.107 | 1, 45 |

**Table S2.** Results of generalized linear models and mixed effects models of ion content relative to water content (WC) in climate-adapted (locational), and seasonally-adapted fly lines. Shaded rows indicate significance based on *P*<0.05.

| Variable | Group | Effect | F | P-value | df |
| --- | --- | --- | --- | --- | --- |
| Na | Locational | WC | 60.33 | <0.001 | 1, 411 |
| Na | Locational | Climate | 0.13 | 0.880 | 2, 12 |
| Na | Locational | Sex | 15.44 | <0.001 | 1, 411 |
| Na | Locational | WC * Climate | 5.59 | 0.004 | 2, 411 |
| Na | Locational | WC * Sex | 0.07 | 0.796 | 1, 411 |
| Na | Locational | Climate * Sex | 2.48 | 0.085 | 2, 411 |
| Na | Locational | WC * Climate * Sex | 4.78 | 0.009 | 2, 411 |
| K | Locational | WC | 579.52 | <0.001 | 1, 411 |
| K | Locational | Climate | 0.59 | 0.570 | 2, 12 |
| K | Locational | Sex | 4.70 | 0.031 | 1, 411 |
| K | Locational | WC * Climate | 25.68 | <0.001 | 2, 411 |
| K | Locational | WC * Sex | 9.72 | 0.002 | 1, 411 |
| K | Locational | Climate * Sex | 0.61 | 0.611 | 2, 411 |
| K | Locational | WC * Climate * Sex | 3.13 | 0.045 | 2, 411 |
| Na | Seasonal | WC | 21.23 | <0.001 | 1, 278 |
| Na | Seasonal | Season | 0.01 | 0.908 | 1, 8 |
| Na | Seasonal | Sex | 30.15 | <0.001 | 1, 278 |
| Na | Seasonal | WC * Season | 0.28 | 0.599 | 1, 278 |
| Na | Seasonal | WC * Sex | 0.08 | 0.784 | 1, 278 |
| Na | Seasonal | Season * Sex | 0.02 | 0.883 | 1, 278 |
| Na | Seasonal | WC * Season * Sex | 3.31 | 0.070 | 1, 278 |
| K | Seasonal | WC | 479.39 | <0.001 | 1, 278 |
| K | Seasonal | Season | 0.44 | 0.528 | 1, 8 |
| K | Seasonal | Sex | 87.68 | <0.001 | 1, 278 |
| K | Seasonal | WC * Season | 5.55 | 0.019 | 1, 278 |
| K | Seasonal | WC * Sex | 0.07 | 0.797 | 1, 278 |
| K | Seasonal | Season * Sex | 12.90 | <0.001 | 1, 278 |
| K | Seasonal | WC * Season * Sex | 10.48 | 0.001 | 1, 278 |

